# Tissue specific dual RNA-seq defines host-parasite interplay in murine visceral leishmaniasis caused by *Leishmania donovani* and *Leishmania infantum*

**DOI:** 10.1101/2022.02.04.479211

**Authors:** Sarah Forrester, Amy Goundry, Bruna Torres Dias, Thyago Leal-Calvo, Milton Ozório Moraes, Paul M. Kaye, Jeremy C. Mottram, Ana Paula C. A. Lima

**Affiliations:** York Biomedical Research Institute, Department of Biology, University of York, York, England, UK; Instituto de Biofisica Carlos Chagas Filho, Universidade Federal do Rio de Janeiro, Rio de Janeiro, Brazil; Instituto Oswaldo Cruz, Fiocruz, Rio de Janeiro, Brazil; York Biomedical Research Institute, Hull York Medical School, University of York, York, England, UK

## Abstract

Visceral leishmaniasis is associated with hepato-splenomegaly and altered immune and haematological parameters in both pre-clinical animal models and humans. We studied mouse experimental visceral leishmaniasis caused by *Leishmania infantum* and *Leishmania donovani* in BALB/c mice using dual RNA-seq to investigate the transcriptional response of host and parasite in liver and spleen. We identified only 4 species-specific parasite expressed genes (SSPEGs; log2FC >1, FDR <0.05) in the infected spleen, and none in the infected liver. For the host transcriptome, we found 789 differentially expressed genes (DEGs; log2FC >1, FDR <0.05) in the spleen that were common to both infections, with IFNγ signaling and complement and coagulation cascade pathways highly enriched, and an additional 286 and 186 DEGs that were selective to *L. donovani* and *L. infantum* infection respectively. Among those, there were network interactions between genes of amino acid metabolism and PPAR signaling in *L. donovani* infection and increased IL1β and positive regulation of fatty acid transport in *L. infantum* infection, although no pathway enrichment was observed. In the liver, there were 1939 DEGs in mice infected with either *L. infantum* or *L. donovani* in comparison to uninfected mice, and the most enriched pathways were IFNγ signaling, neutrophil mediated immunity, complement and coagulation, cytokine-chemokine responses and hemostasis. Additionally, 221 DEGs were selective in *L. donovani* and 429 DEGs in *L. infantum* infections. These data show that the host response for these two visceral leishmaniasis infection models is broadly similar, and ∼10% of host DEGs vary in infections with either parasite species.

**Importance:** Visceral leishmaniasis (VL) is caused by two species of *Leishmania* parasites, *L. donovani* in the Old World and *L. infantum* in the New World and countries bordering the Mediterranean. Although cardinal features such as hepato-splenomegaly and alterations in blood and immune function are evident, clinical presentation may vary by geography, with for example severe bleeding often associated with VL in Brazil. Although animal models of both *L. donovani* and *L. infantum* have been widely used to study disease pathogenesis, a direct side-by-side comparison of how these parasites species impact the infected host and/or how they might respond to the stresses of mammalian infection has not been previously reported. Identifying common and distinct pathways to pathogenesis will be important to ensure that new therapeutic or prophylactic approaches will be applicable across all forms of VL.

## Introduction

Visceral leishmaniasis (VL) is responsible for 50,000-90,000 new cases a year, and the majority (>94%) of reported new cases occurs in seven countries. It is caused by two parasite species, *L. donovani*, which is associated with infections mainly in Africa and in Southern Asia, and *L. infantum*, which is associated with infections in South America, Middle East and Southern Europe. Both host and parasite genetics contribute to the manifestation of disease in infected individuals, and factors such as the *Leishmania* species, health status of the host, and urbanisation may contribute to the disease progression and pathogenesis. VL is lethal if left untreated, and is often associated with anaemia, thrombocytopenia, hepatosplenomegaly and downregulation of immunity in both humans (1, 2) and pre-clinical rodent models (3, 4). There are significant differences in the pathogenesis of human VL between India, East Africa and Brazil. Lymphadenopathy is common in East Africa but rare in India; post-kala azar dermal leishmaniasis (PKDL) is frequent in East Africa and less so in India but does not occur in Brazil. On the other hand, severe bleeding, and coagulopathy, often linked to disseminated intravascular coagulation (DIC) has been reported in VL in Brazil, where it is a major risk factor for death, whereas in India and Sudan, bleeding is most commonly limited to epistaxis. The severity of VL in patients from Brazil, including coagulopathies, has been associated with a higher concentration of serum inflammatory cytokines, although this is ineffective in the control of parasite proliferation (5, 6). A study of parasite burden in patients infected with *L. infantum* revealed a direct correlation between burden and clinical outcome, disease severity and mortality risk (7, 8). How and which parasite factors influence the outcome of infection and the development of distinct clinical conditions, however, remains elusive.

In the last few years, attempts to identify blood signatures in infected patients using transcriptomics helped to define general pathways involved in the host response of symptomatic individuals, such as T-lymphocyte activation and type I-interferon signalling in VL patients infected with *L. infantum* (9), or enrichment of genes associated with erythrocyte function in active cases of VL patients infected with *L. donovani* (9). One transcriptomics approach is dual sequencing, which allows for both pathogen and host responses to be characterised simultaneously within the same biological sample and can enable a greater understanding of the host-pathogen interactions. Dual RNA-seq has been used to investigate gene expression changes in *in vivo* models of *Mycobacterium tuberculosis* infected murine lungs (10), infection of the fish *Epinephelus coioides* with *Pseudomonas plecoglossicida (11)*, and neutrophil recruitment in *Streptococcus pneumoniae* murine infections (12).

In the study of leishmaniasis, dual RNA-seq has been used to analyse the parasite-macrophage interaction *in vitro* (13–15), and in human lesion material from *L. braziliensis*-infected patients (16). In contrast, dual RNA-seq has not to date been applied to the study of visceral leishmaniasis in animal models. In addition, whilst previous studies have examined the host transcriptional response to *L. donovani* (17) and *L. infantum* (18) infection in mice, as well as hamsters infected with *L. donovani (19)*, direct comparisons of the host response between these two visceralizing species has not been reported previously. Likewise, although both *L. donovani* (20) and *L. infantum* (21) have been well characterised in terms of gene expression related to stage specific differentiation, it is not known if these two parasites react similarly to the stresses associated with mammalian infection.

Given the potential for distinct pathologies to be provoked by *L. donovani* and *L. infantum* infection and the potential diversity of tissue gene expression and tissue-specific immunopathology, we aimed to investigate simultaneously the host and parasite global gene expression changes within key target organs in infected mice. We performed dual RNA-seq analysis to compile a complete list of differentially expressed host genes (DEGs) as well as the species-specific parasite expressed genes (SSPEG). Our data indicate extensive similarity in host response, but with some infection-specific differences. In contrast, *L. donovani* and *L. infantum* isolated from spleen and liver show very few differences in gene expression.

## Results

### Parasite transcriptional profile during *L. donovani* and *L. infantum* infections

We examined host and parasite response at day 35 post infection in BALB/c mice, as this time point reflects both the establishment of chronic splenic pathology and the onset of hepatic host resistance (17). We applied tissue dual RNA-seq to liver and spleen tissue isolated from uninfected mice and mice infected with either *L. donovani* or *L. infantum* amastigotes (**Fig. 1A**). Spleen and liver parasite burdens (as assessed by mRNA abundance for *Amastin*) are shown in **Fig. 1B**. The extent of hepato-splenomegaly is shown in **Fig. 1C** and was not significantly different between the two infections.

**Fig. 1.**
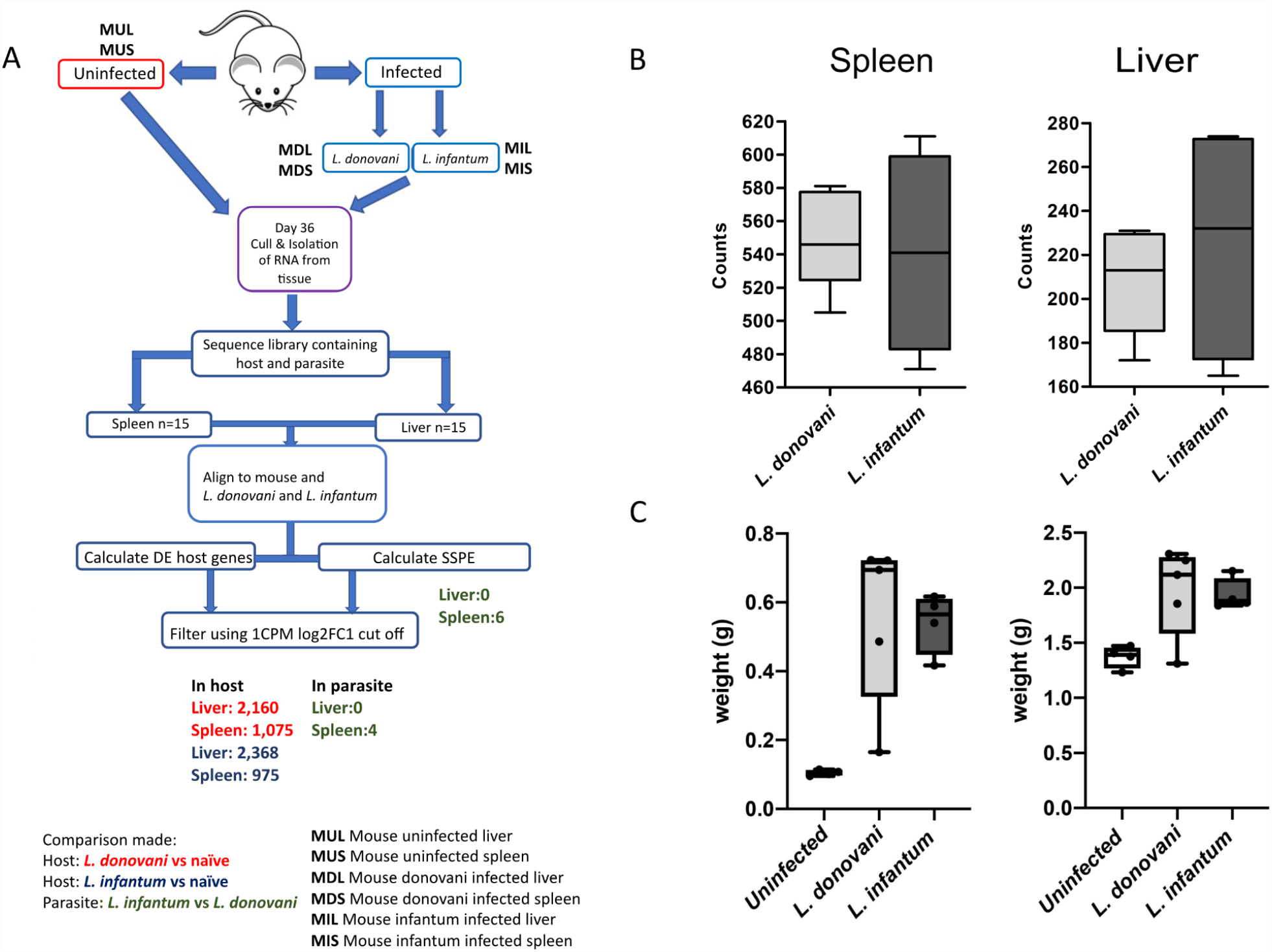
Dual-RNASeq analysis of *L. donovani* and *L. infantum* infection in BALB/c mice. **(A)** Schematic representation of the experimental design. BALB/c mice were infected for 35 days (n=5 mice per group). Abbreviations; M1-5, mouse 1-5; U, uninfected; S, spleen, L, liver; Mouse 1 uninfected spleen: M1US. Numbers shown represent DE genes. **(B)** Amastin counts. Boxplot showing the normalised read count distribution for spleen and liver of infected animals based on the amastigote-expressed gene amastin LdBPK_080710.1 (for *L. donovani*) and ortholog LINF_080012100 (for *L. infantum*) **(C)** Organ weights for spleen (left) and liver (right) in uninfected and infected animals. Box (25^th^ to 75^th^ percentiles) and whisker (min, max) plots with median (line). No significant differences were observed (Mann-Whitney, *p*-value > 0.05).

For those genes that have syntenic orthologues in both *L. donovani* and *L. infantum*, we identified 7250 *L. donovani* and 7227 *L. infantum* genes expressed in the spleen of an infected mouse and 7092 *L. donovani* and 7070 *L. infantum* expressed genes in the liver (**Table S1** in the supplementary material). In both the spleen and liver, the 100 most abundant mRNAs in both species were mainly involved with parasite metabolism. These represent 35 GO term Biological function pathways, with 53 of the genes being ribosomal proteins and therefore associated with translation (GO:0006412). **Fig. 2** shows the top 50 most abundant mRNAs excluding ribosomal genes, to emphasise additional biological processes which include heat shock proteins, tubulin, histones and ubiquitin. Of note, six amastin glycoprotein mRNAs (*ama8B, ama34D* and *ama34A* families) were among the most abundant, in keeping with the importance of these surface proteins to tissue amastigotes (22). mRNAs for six hypothetical proteins of unknown function, but conserved in trypanosomatids, were also highly abundant, as was mRNA for the stage-regulated lysosomal cathepsin cysteine peptidase (CPB), a known virulence factor (23). The expression of tryparedoxin peroxidase, an enzyme crucial for detoxification (24), was also among the 50 top expressed genes by amastigotes of both species, in both spleen and liver.

**Fig. 2.**
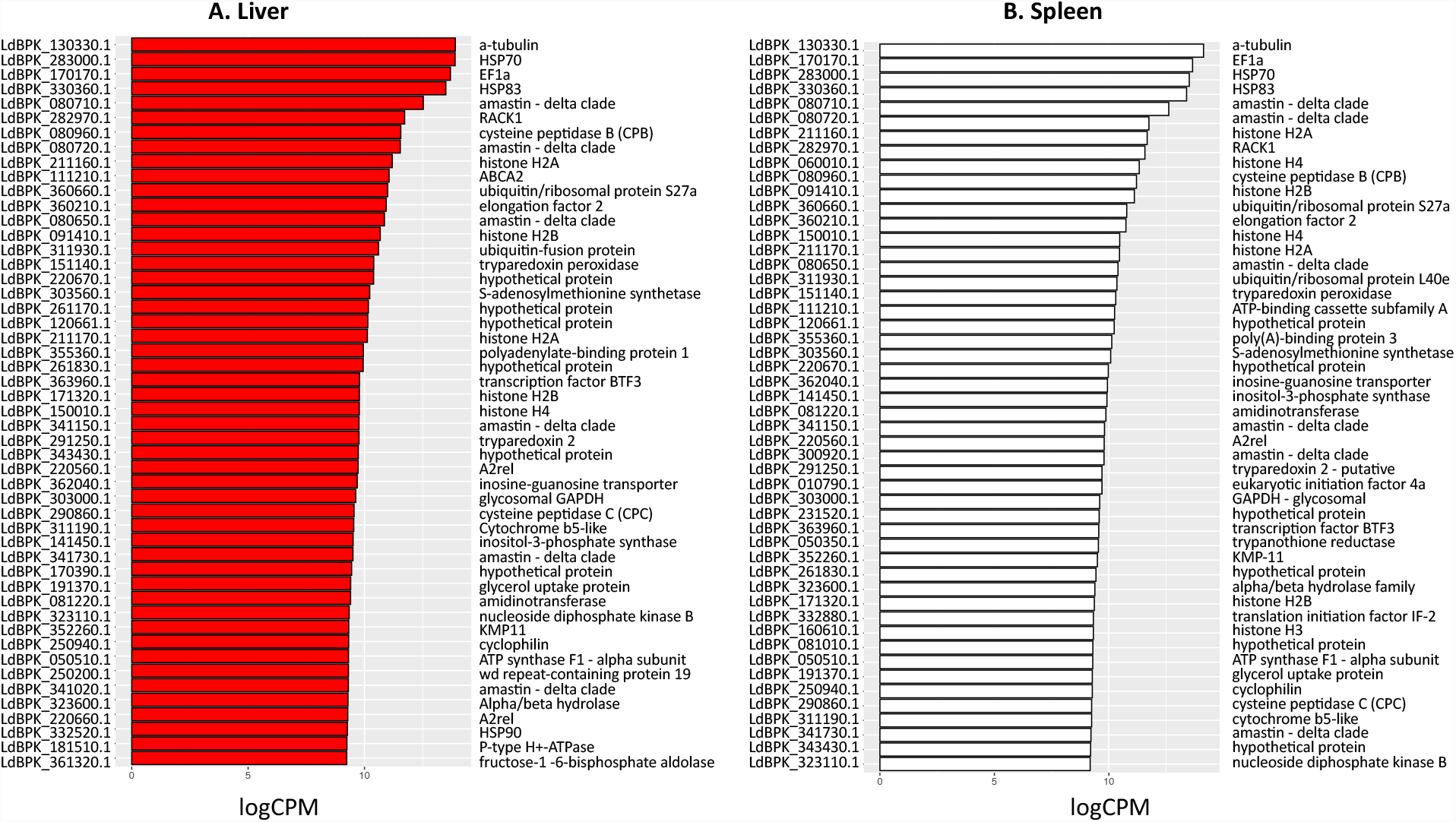
The 50 most highly expressed parasite genes in the liver **(A)** and spleen **(B)** excluding ribosomal proteins based on logCPM values shown in **Table S1**. *L. donovani* gene annotations are used here, the syntenic *L. infantum* orthologues gene IDs can be found in **Table S1**.

### Species-specific parasite expressed genes

We then asked if there were species-specific parasite expressed genes (SSPEGs) between different organs of infected mice. To address this, we used a combined minimum of ten *L. infantum* and *L. donovani* transcripts from all nine animals as a cut off for the expression analysis. 6,947 genes passed this filter in the spleen and 6,285 in the liver (**Fig. 3** and **Table S1**). In the liver, 119 genes had zero counts in *L. donovani* but were expressed in *L. infantum*. Among those, there were four genes encoding ABC proteins, six protein kinase genes, and enzymes involved in detoxification, such as glutaredoxin and glutathione peroxidase. 141 genes had zero counts in *L. infantum* but were expressed in *L. donovani* in the liver (**Table S1)**. Among those, there were protein kinases, ubiquitin-associated enzymes, and a pentamidine-resistance protein. In the spleen, 57 genes had zero counts in *L. donovani* but were detected in *L. infantum*, while 80 genes had zero counts in *L. infantum* but were detected in *L. donovani*, including glutaredoxin (**Table S1**). As shown in Fig. 1A, there were only 4 SSPEGs following filtering in the spleen, but none in the liver (**Fig. 3C and D** and **Table S1**). Within the 4 SSPEGs, the Lsmp7 protein gene (LINF_260005000) is expressed by *L. infantum* in the spleen of infected mice, but there were zero counts in *L. donovani*. Three hypothetical proteins (LdBPK010410, LdBPK040220 and LdBPK231520) were more expressed by *L. donovani* (log_2_FC>1.3 in comparison to *L. infantum*); while the first gene is conserved among trypanosomatids, the last two are absent from *Trypanosoma* species and could be associated with *Leishmania*-specific molecular pathways.

**Fig. 3.**
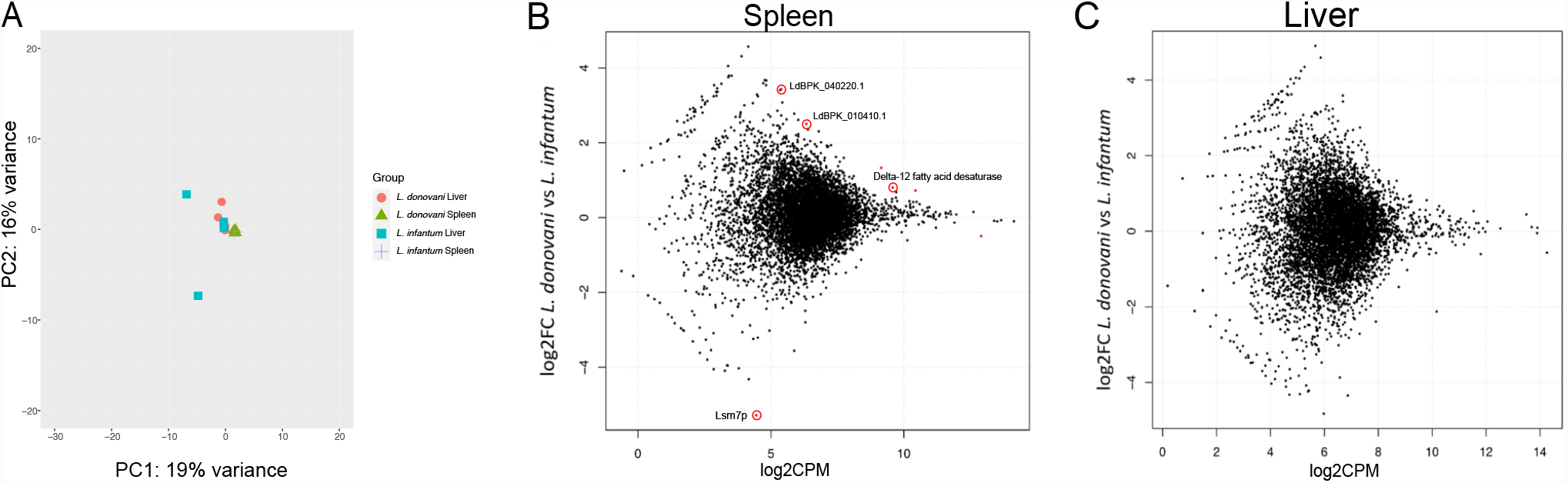
**(A)** Principal component analysis of the parasite transcriptome for the spleen and liver. MA plots of SSPE genes between *L. donovani* and *L. infantum* in spleen **(B)** and liver **(C)**. MA plots show the logged ratio between read counts averaged per group, per gene. Genes with an FDR value of < 0.05 are shown in red. MA plots for liver vs spleen in *L. infantum* infections and *L. donovani* infections are shown in Fig. S1.

Due to low levels of variation between groups, there was no distinction in parasite gene expression between *L. infantum* and *L. donovani* in liver or spleen, as determined by Principal Component (PCA) analysis (**Fig. 3A**). The fact that we could not identify SSEPGs in the liver is not due to lack of read depth, as a high number of expressed parasite genes were identified (**Table S1**). The differences in gene expression between the *Leishmania* species are, therefore, tissue specific. Paralog counts for each of the 4 SSPEGs were calculated in both the *L. donovani* BPK208 and *L. infantum* JPCM5 reference genomes to check whether the significant differences in mRNA abundance between the two species found in the spleen was due to differences in gene copy number between species. Three of the four SPPEGs had the same number of paralogs, and the delta-12 fatty acid desaturase had one additional paralog in *L. infantum* **(Table S2)**. The identification of differential expression (DE) genes was also done using the transcript per million method, which controls for gene length, in order to identify false positive SPPEGs due to orthologue length differences. However, this did not result in additional differentially expressed genes. In addition, reads per kilobase of transcript per million mapped reads (RPKM) values were also generated and correlated between log2FC RPKM and log2FC counts per million (CPM) values, which had a R squared of 0.93 using a filter of log2FC of between -1 and 1. Of all expressed parasite genes, only 48 genes didn ‘t correlate, and had a positive log2FC RPKM value and a negative log2FC CPM value. The length of these orthologues in the *L. infantum* reference and the length of the orthologue in the *L. donovani* reference for these genes were plotted (**Fig. S1A)**, and 45 of these genes had an identical gene length between the orthologues. This suggests that gene length differences between orthologues does not result in either false positive or false negative SPPEGs.

### Differences in the *Leishmania* transcriptome is neither tissue nor species dependent

Gene expression was determined for parasite transcripts in both the spleen and liver for each parasite species to observe whether any differential expression was observed when parasites infect different tissues (**Fig. 3C and Fig. S1B**,**C**). In both *L. donovani* and *L. infantum* infections, PCA analysis showed no separate clusters for liver and spleen samples for the parasite transcriptome (**Fig. 3A**). It also shows that there are multiple components with small contributions to the variation, with only 35% explained by the first and second components. Only one parasite gene, histone H4, was DE in *L. donovani* when comparing spleen and liver, and none in *L. infantum*. These analyses indicates that there is no detectable discriminatory transcriptional signature between spleen and liver amastigotes for either *L. donovani* or *L. infantum* infections.

### Host transcriptional response to *L. donovani* and *L. infantum* infection

The host response to infection with *L. donovani* in murine (17, 25) or hamster models (19, 26), as well as murine infection with *L. infantum* (21) have previously been described, but a direct comparative study has not previously been undertaken of these two parasite species in the same host genetic background. BALB/c mice infected for 35 days had similar parasite burdens in spleen and liver as assessed by read counts for *Amastin* (LdBPK_080710.1 in the *L. donovani* BPK reference and *L. infantum* orthologue LINF_080012100; **Fig. 1B**) and similar degrees of hepatosplenomegaly, as assessed by organ weight (**Fig. 1C**).

PCA analysis shows that there is clustering based on infection status, as well as on the organ specific host response, with 99% of the variance explained along PC1, where liver and spleen samples cluster separately (**Fig. 4A**). In comparison, there is infection status clustering, however this variance is much lower compared to the organ specific differences observed, as shown by the 1% variance explained by PC2. This variance between uninfected and infected individuals is larger in the liver compared to the spleen (**Fig. 4Bi** and **Fig. 4Bii**, and the clustering in **Fig. 4A**). In *L. donovani* infections, we identified 2,160 DE host genes in the liver and 1,075 in the spleen relative to uninfected controls (**Fig. 4B** and **Fig. S2A-D**). DE genes were identified with a cut off of log2FC > 1, CPM > 1 and false discovery rate (FDR) value of < 0.05. In *L. infantum* infections, there were 2,368 DE genes in the liver and 975 in the spleen, relative to uninfected controls. 1,939 DEGs were commonly identified across both infections in the liver, representing 89.8% and 81.9% of all infection induced genes in *L. donovani* and *L. infantum* infected mice respectively. There was 93% correlation in the liver and 94% in the spleen between the uninfected and infected log2FC in the *L. donovani* and *L. infantum* infections (**Figs. 4C, D and Figs. S2E,F**). The greatest outlier, shown in both A and B is Nxph4. However, this is due to >700 counts in two individuals versus less than five in the remaining 3 individuals. The other outlier is Coch, however this is not significantly different between *L. infantum* and *L. donovani* (**Table S3**). In spleen, 789 DEGS were common between the two infections, representing 73.4% and 80.9% of all infection induced genes in *L. donovani* and *L. infantum* infected mice respectively (**Fig. 4C** and **Table S3**). In addition to these commonly expressed genes, the livers of mice infected with *L. donovani* and *L. infantum* had 221 and 429 unique DEGs, and the spleen had 286 and 186 unique DEGs for *L. donovani* and *L. infantum*, respectively.

**Fig. 4.**
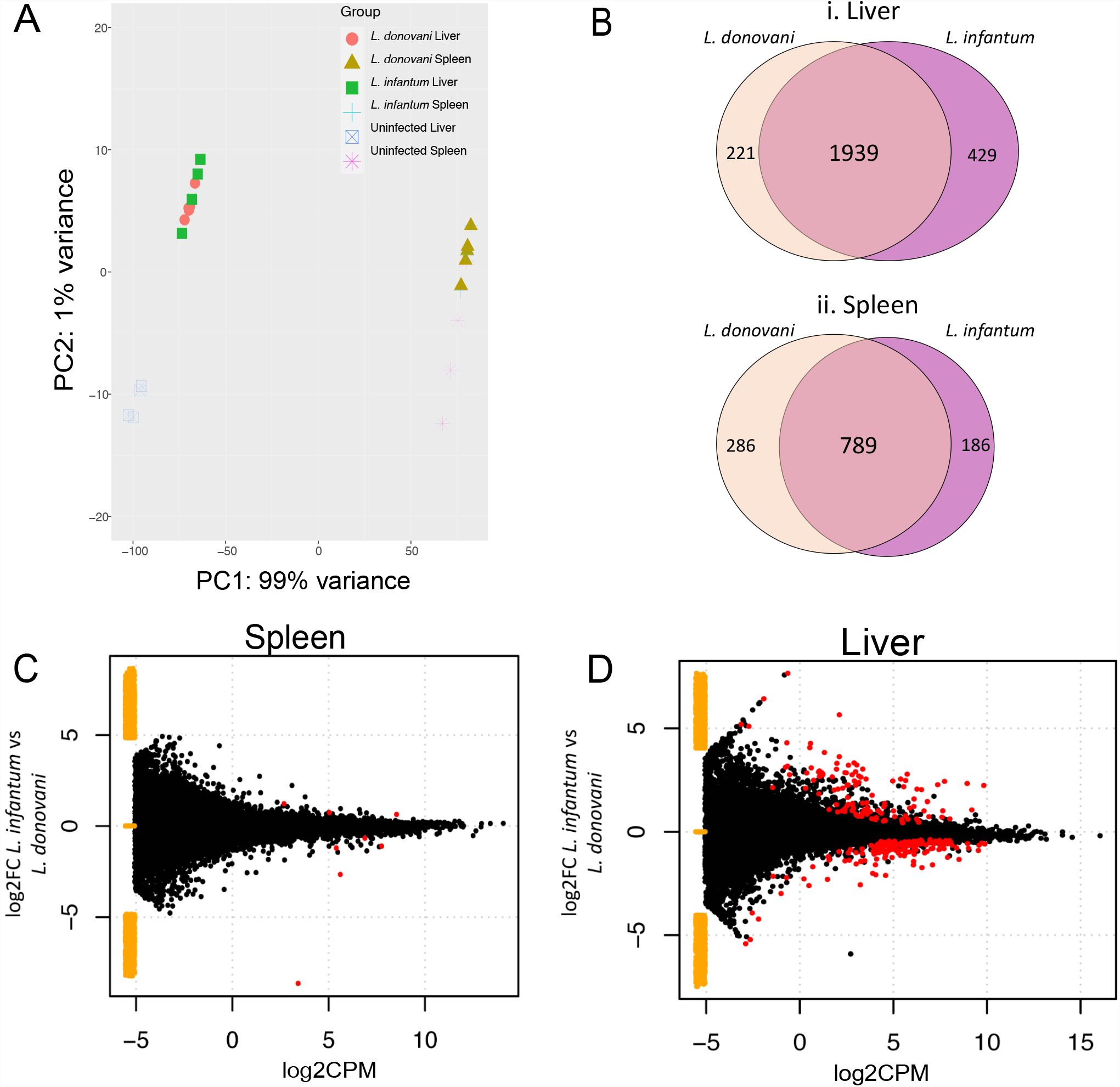
**(A)** PCA analysis based on the host genes for the spleen and liver for *L. donovani* infected, *L. infantum* infected and uninfected samples. Comparisons between uninfected and *L. infantum* infected and uninfected and *L. donovani* infected mice in the liver and spleen are shown in **Fig. S2**. Venn diagram of host genes in the liver **(B) i** and in spleen **(B) ii**. Overlap shows the parasite genes that are DE in both infections. In the liver there were 2160 DE host genes in *L. donovani* and 2368 host DE genes for *L. infantum*. In the spleen, there were 1075 host DE genes for *L. donovani* and 975 host DE for *L. infantum*. MA plot with the DE genes identified in host genes between *L. infantum* and *L. donovani* infected mice in the spleen **(C)** and in the liver **(D)**. Yellow points represent genes expressed in one condition, but not the other.

As an example of potentially relevant unique DE genes, arachidonate 5-lipoxygenase expression (Alox5) was increased solely in the *L. donovani*-infected spleen (log2FC 1.09). Since this enzyme is a major player in the synthesis of leukotrienes and arachidonic-acid, it is possible that such lipid mediators are also increased in the *L. donovani* spleen (27, 28). Due to its capability to metabolize oxidised fatty acids and generate lipoxin and resolvins as well, it can also contribute to counteract inflammation (28, 29). Since RNA-seq studies cannot give a picture of the relative amounts of lipid mediators in the tissue to allow predictions on the degree of inflammation, metabolomic studies would be more appropriate to address such potential differences (30). Among the unique host DE genes for each parasite species, some were related to the immune response. For example, we detected significant increased expression of the chemokine receptor CCR2, and of matrix metalloprotease (MMP9), and significant decreased expression of CCR4 and CXCR5 in *L. donovani* infected spleens, while the levels in *L. infantum* infected spleens were not significantly different from the uninfected spleen. In *L. donovani* infected spleens, TNFα upregulation also passed the filter as significantly different from uninfected controls, while not in the *L. infantum* infected spleen. Likewise, among unique DE genes, there was a significant decrease in the expression of CCR9 in the spleen and increase of CXCR5 in the liver of *L. infantum* infected mice. Chemokine receptor expression can have a great impact in the homing of immune cells. Since we cannot predict if the changes in the expression for chemokine receptors occur in specific cell types, it is possible that greater differences of host species specific DE genes will be observed when RNA-seq of specific cellular subsets purified from the infected spleen is performed. This might be relevant in the context of cellular recruitment to tissues, since the protein gradient of chemokines laid on the extracellular matrix combined with the amount of surface expression of their receptors in immune cells dictate the timing, specific cellular subtypes and number of cells that enter the infected organs. More strikingly, the expression of IL1β, a cytokine that results from inflammasome activation, was significantly increased only in the *L. infantum*-infected spleen. Generation of bioactive IL1β depends on caspase 1 activation, which was expressed in the infected spleen, creating a scenario where IL1β-dependent downstream effects could take place in *L. infantum* infection. On the other hand, the expression of an inflammasome gene, NLRP3, appeared as specifically upregulated in the *L. donovani*-infected spleen, while no elevation in IL1β expression was observed. In agreement with that, it has been proposed that in infected macrophages, *L. donovani* subverts inflammasome activation and the consequent cellular pyroptosis (31).

To obtain a broad overview of immune related pathways, we examined cytokine, chemokine and TLR genes that were significantly DE in infected mice (**Fig. 5**). When analysed as a whole, the host response to *L. donovani* and *L. infantum* infections for lymphokines and immune response-related enzymes is largely similar in both organs. Many of the host DE genes common to both infections correspond with pathways associated with a generalised immune response, as described previously for *L. donovani* (17). Analysis of DE genes common to both *L. donovani* and *L. infantum* vs naïve in the liver shows that the GO:BP immune system process is the most significantly enriched pathway (adjusted p-value 3.45×10^−110^) followed by immune response (adjusted p-value 2.634× 10^−85^). The five most enriched GO term BP annotations in the liver correspond with immune system related gene expression (**Fig. S3A, S3B**). The five most enriched annotations in the spleen are the same as the most enriched in the liver. In addition, the response to stress is also amongst the top annotations in the spleen (**Fig. S3C, S3D**). The analysis of BALB/c mice infected with *L. donovani* (17), also showed the extent of the immune response related DE genes is greater in the liver than in the spleen. This is also suggested in this dataset by the enrichment in the top GO BP term for both organs, Immune system process, which has an adjusted p-value of 3.45×10^−110^ in the liver, but 1.034×10^−81^ in the spleen. Among those, the highest fold change in the infected liver were for *Nos*2 (log2FC>10), *Ifn*γ (log2FC>8), *Tnf* (log2FC>5), and *Il21* (log2FC>6.6), denoting the sustained high inflammatory environment in this tissue during late infection. In addition, we observed high levels of expression of *Il*27 (log2FC>4.8), which was not previously observed as a DE gene in micro-array studies (17).

**Fig. 5.**
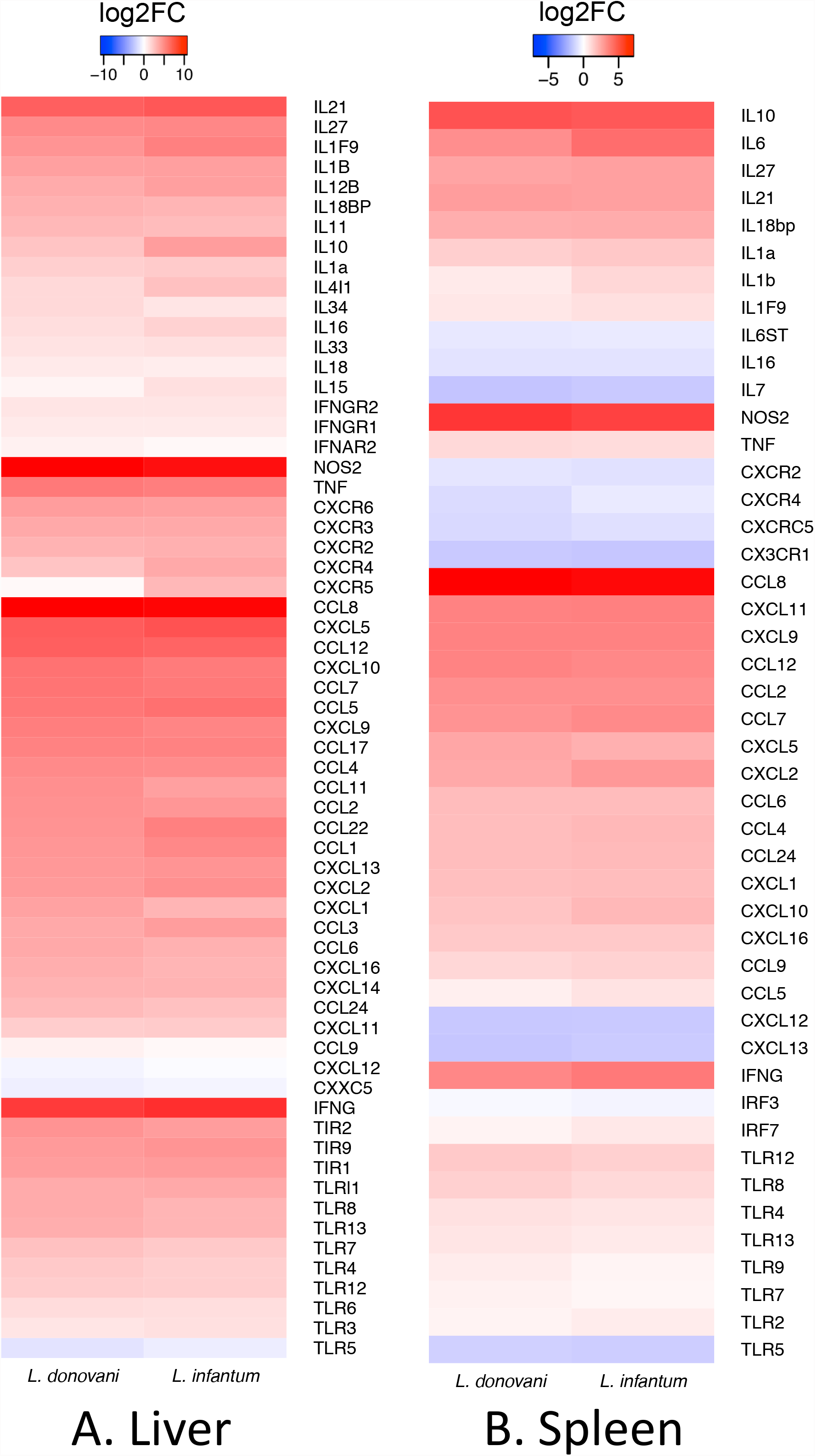
The immune response-related DEGs of the host in *L. donovani* and *L. infantum* infections in the liver **(A)** and spleen **(B)**. The colours represent averaged log2FC changes per group. Red represents genes that are increased in expression following infection, and blue are decreased following infection, in comparison to uninfected controls.

In the spleen, *Nos*2 (log2FC>5) has the highest fold change in mRNA abundance amongst defence-related responses, while *Tnf* (>0.95 log2FC <1.05) and *Ifn*γ (log2FC3) had lower fold change increases in mRNA abundance compared to that seen in liver (**Fig. 5**). Unlike in the liver, where *Il6* mRNA abundance was not increased during infection, *Il6* mRNA increased in abundance by >8 fold in the spleen. *Il10* was DE in both *L. donovani* and *L. infantum* in the spleen and liver. In the spleen the log2FC>4.5 in both infections relative to naive, whereas in the liver the log2FC is lower in naïve vs *L. donovani* infected (log2FC 2.5) compared to *L. infantum* infected (log2FC 4).

Despite a generalised common defence pathway being observed in both tissues for both parasite species, it is possible that the small, but significantly different, changes in the expression of immune response related genes found in the tissues infected with *L. donovani* versus *L. infantum* could combined, contribute synergistically to provide substantially distinct micro-environments with consequences for long-term pathology.

Next, gene enrichment analysis was performed on the common gene set that were DE in infected mice as compared to uninfected controls using EnrichR and significantly enriched pathways were identified within GO Biological Processes, GO Molecular Function, Wiki Pathways and KEGG. In the liver (**Table S4**), the most significantly enriched terms included those related to IFNγ signalling (inflammatory response), neutrophil mediated immunity, coagulation pathways and hemostasis. A similar analysis was also performed on the 221 genes identified as DE only in *L. donovani*-infected mice in the liver and no enriched pathways were identified. The 429 mouse genes only DE in *L. infantum* infected mice in the liver showed significant enrichment in genes associated with glycine, serine and threonine metabolism (adjusted *p*-value 1.34E^-07^) with cysteine and methionine metabolism (adjusted *p*-value 1.02E-^06^) and in cell cycle related genes. An overview of the enriched pathways identified from DE common to both species is shown in **Fig. 6**.

**Fig. 6.**
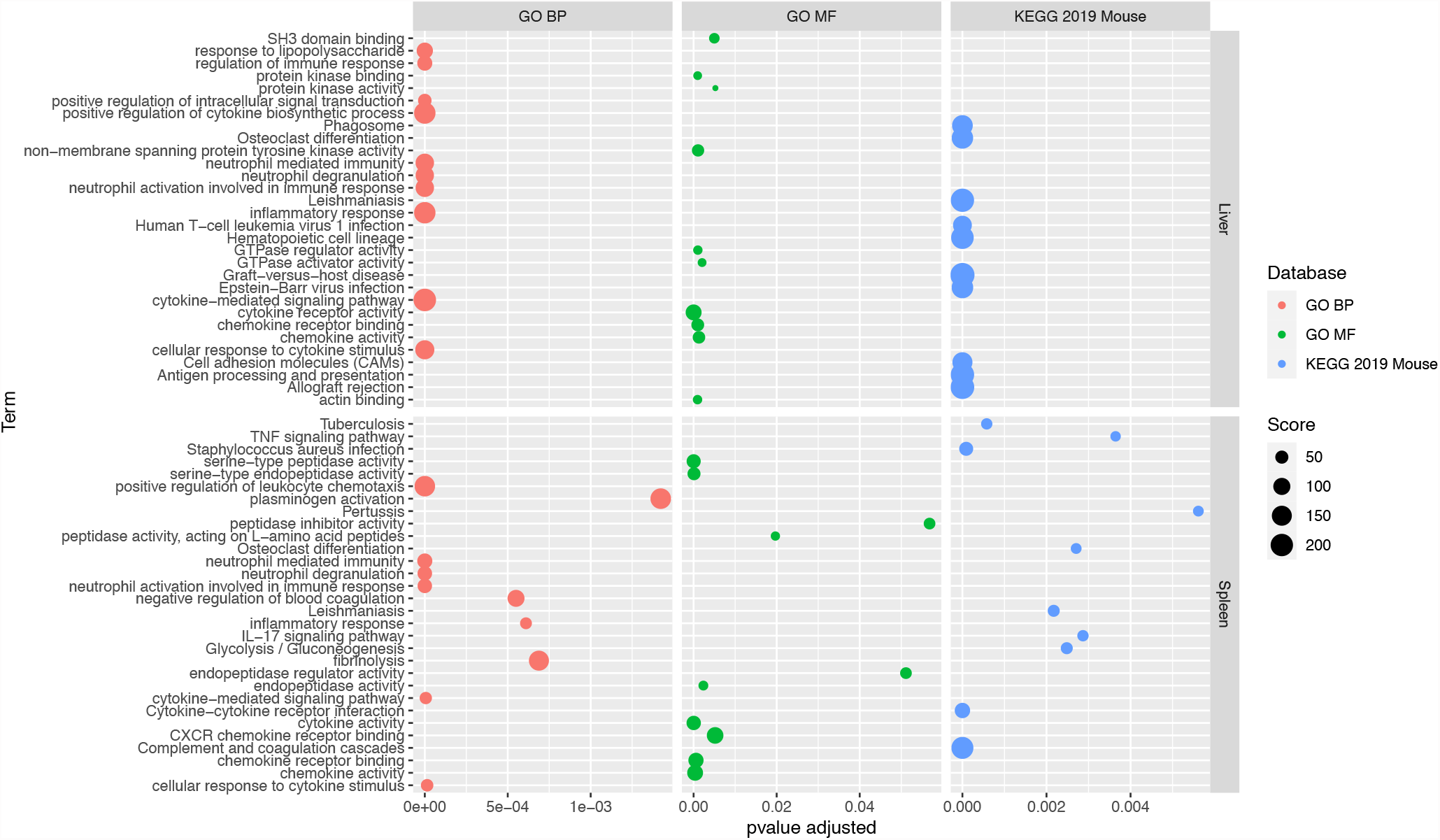
Summary plot of EnrichR data in **Tables S4 and S5**. The adjusted p-values (Benjamin Hochberg correction) showing the significance of the enrichment of the term are shown on the x axis. The size of the dot represents the score of the enrichment based on significance and the number of genes involved in the enrichment, with higher scoring enrichments being more significant. The databases used are Gene ontology biological processes (GO BP), Gene ontology molecular processes (GO MF) and Kyoto Encyclopaedia of Genes and Genomes Mouse specific database (KEGG 2019 Mouse).

In the spleen, enriched terms common to both infections mirrored those in the liver (**Table S5)**. Immunity and chemotaxis were highly enriched, with additional prevalence of hemostasis-related biological processes in both directions, such as clotting cascade, plasminogen activation and platelet degranulation, as well as negative regulation of blood coagulation and fibrinolysis. Network analysis of the 286 DE specific to *L. donovani* infection in the spleen showed that pathways related to phenylalanine metabolism and phenylalanine, tyrosine and tryptophan biosynthesis were significantly enriched (adjusted *p*-value 0.044 and 0.047 respectively). Tryptophan levels have been associated with leishmaniasis, by means of its regulation by the rate limiting enzyme indoleamine 2,3-dioxygenase (IDO-1). IDO-1 also plays a role in immune regulation as a molecular switch, mainly by its early expression in antigen presenting cells, that leads to reduced levels of tryptophan in the surrounding micro-environment, with deleterious consequences to T-cell activation (32). IDO-1 inhibitors were shown to reduce parasite burden and pathology (33), and it was proposed as a marker for immunosuppression in VL (34). IDO-1 was also described as required for successful infection of mice with *L. major* or *Toxoplasma gondii* (35) and enhanced expression of IDO-1 was highly discriminatory in lesions of leprosy patients (36). Considering that IDO-1 is upregulated in the *L. donovani* infected spleen (log2FC=2.2; FDR <0.011), together with enhanced *tat* expression, an enzyme that catalyses conversion of tyrosine, it is plausible that a change in nutritional metabolism occurs more evidently in *L. donovani* infections, while not observed in the *L. infantum* infected mice. There was no significant enrichment in the 186 genes DE only in *L. infantum* infections, however it is noteworthy that chemokine receptors CCR9 and CCRL2 are among those DEGs.

Collectively, these data suggest significant overlap in host response to these two viscerotropic species of *Leishmania*, commensurate with similar parasite burdens and gross tissue pathology.

### Protein networks of differentially expressed host genes inferred from Pathway Analysis

Protein interaction networks were predicted from the host genes that were differentially expressed using STRING (**Fig. 7 and Table S6**). STRING predicted a medium sized network from the *L. donovani* specific DE genes in the spleen. Within this network were three significantly enriched KEGG pathways, which correspond with GO term enrichment. These were PPAR signalling, phenylalanine, tyrosine and tryptophan biosynthesis and phenylalanine metabolism, which all have an adjusted *p*-value of 0.0244 (**Fig. 7A**). Nevertheless, cytokine production and fatty acid transport pathways were still significant (*p*-adjusted values of 0.0311 and 0.0257 respectively) (**Fig. 7B**).

**Fig. 7.**
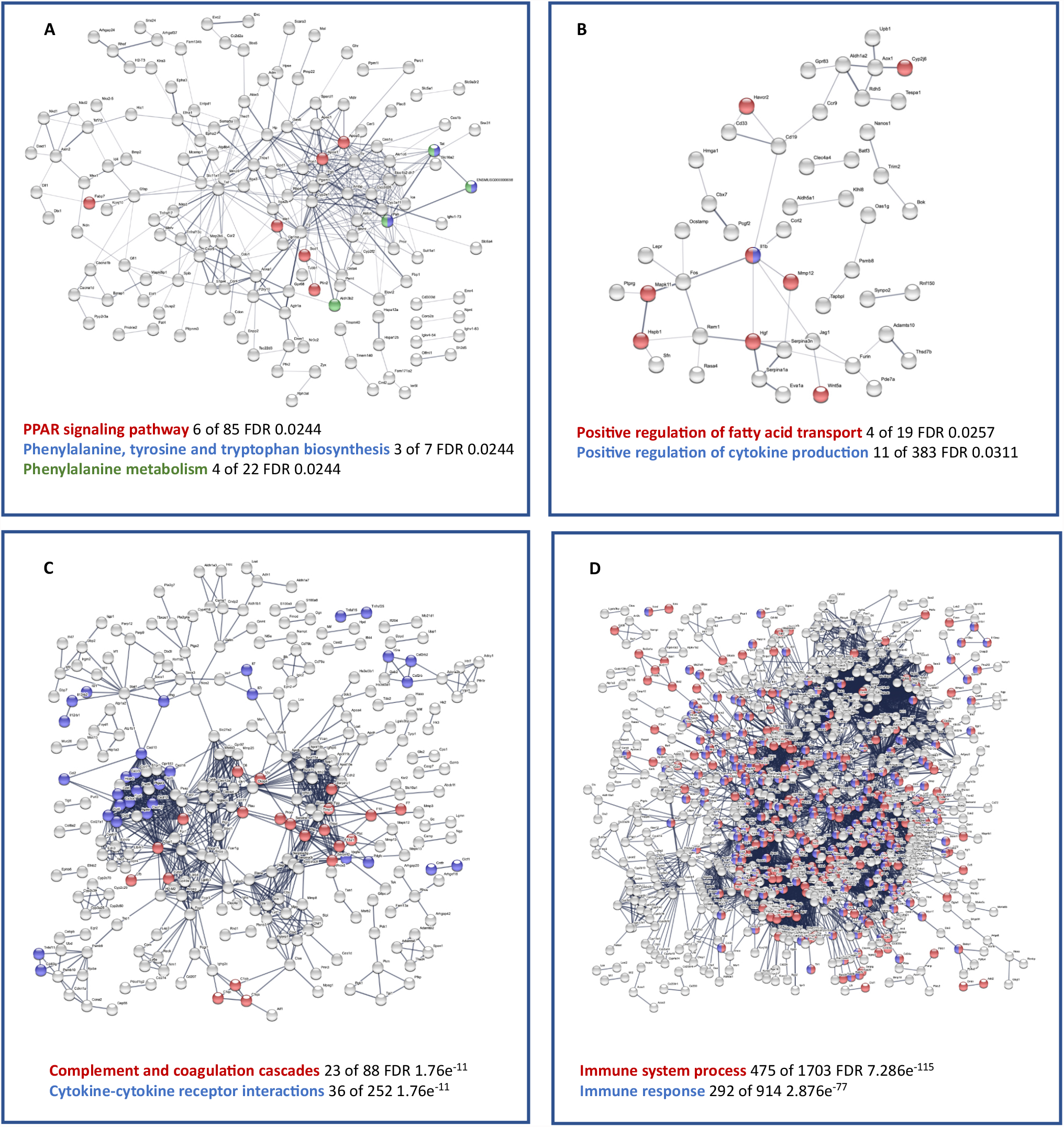
Network analysis was performed using STRINGDB annotations in the spleen using host genes. **(A)** shows interactions between proteins encoded by genes that are specifically DE in *L. donovani* infected vs uninfected in the spleen. **(B)** shows the network of DE genes that are specifically DE in *L. infantum* infected in the spleen. **(C)** shows the network of genes DE in the spleen which are common to *L. infantum* or *L. donovani* infected mice vs naïve. **(D)** Shows the network of genes DE in the liver which are common to *L. infantum* or *L. donovani* infected mice vs naive. Genes that are disconnected nodes are not shown in the Figure but are included within the enrichments highlighted for each panel. Genes within these enrichments are given in **Table S6**.

Within the *L. donovani* infected liver DE specific genes also result in a medium sized network and has significantly enriched interactions within the network (PPI enrichment *p*-value 1.61e^- 07^), however, these are also largely representative of base metabolic processes such as regulation of biological process (FDR 2.86e^-06^) and response to stimulus (FDR 5.84e^-07^). *L. infantum* DE specific genes also predict a significantly enriched network (PPI enrichment p-value 6.11e^-15^) based on non-specific metabolic terms such a metabolic process (FDR 2.32e^-12^) and primary metabolic process (FDR 2.43e^-11^).

DE genes common to both infections in the spleen resulted in a large and robust network (PPI enrichment < 1.0e-^16^). A highest confidence score (0.9) version of this network is shown in **Fig. 7C**. Enrichments were identified in the innate immune system, both Reactome pathways (FDR 4.3e-^13^) and complement and coagulation cascades (FDR 1.76e^-11^). Cytokine-cytokine receptor interactions were also highly enriched (FDR 1.76e^-11^). In the liver, DE genes common to both infections resulted in a significant network (PPI enrichment p-value < 1.0e^-16^), however these genes correspond largely with a generalised immune response, with immune system process and immune response amongst the most significant pathways (FDR 195e-^97^ and 2.21e^- 62^ respectively), see **Fig. 7D**.

### Systemic signature associated with VL

In a micro-array analysis of BALB/c mice infected with *L. donovani*, we previously identified a 26 gene signature that was DE between uninfected and infected mice at days 15,17, 36 and 42 post infection in blood, spleen and liver (17). In the current study, we also observed that 25 genes from the 26 gene signature were also DE relative to uninfected mice in spleen and liver (**Fig. S4**). *Gbp1* does not have an ENSEMBL ID for the *Mus musculus* annotation that corresponds between the database used in this analysis and the version used in Ashwin et al., 2019. *Irg1* is represented by *Acod1* in both datasets and labelled as such. Hence, this systemic signature not surprisingly is also observed in *L. infantum* infected mice.

## Discussion

We used dual RNA-seq to address the following main questions in experimental VL (i) are there any transcriptional differences between the parasites, *L. infantum* and *L. donovani* in different tissues and (ii) what are the similarities and differences in the liver and spleen between infection with *L. infantum* and *L. donovani* compared to uninfected animals. We compiled a complete list of expressed genes for two different visceralising parasite species, *L. donovani* and *L. infantum*, and two infected organs, the liver and spleen, at a single time-point of infection. This time-point was chosen to reflect both the establishment of chronic splenic pathology and the onset of hepatic host resistance, from the commonly used BALB/c mouse model.

The identification of genes expressed by amastigotes in the parasitised tissues in chronic infections could help to narrow down the panel of genes within their genomes when looking for potential virulence factors and/or drug targets. Surprisingly, the sets of expressed genes were nearly identical in the two parasite species, collected from the two organs. Considering that in the mouse model, the infection is contained and resolved in the liver while parasite burdens increase in the spleen, one might have expected that different sets of expressed genes would be mobilised by the parasite to respond to those differing tissue microenvironments. In *Leishmania*, however, gene-expression regulation can also occur at the post-transcriptional level making it possible that tissue-specific signatures of parasite genes would be preferentially identified in proteomics studies.

Previously validated virulence factors such as cysteine peptidase B (CPB) and tryparedoxin 1 were among the top 50 most highly abundant mRNAs in the liver, with cysteine peptidase C (cathepsin B-like peptidase) and cysteine peptidase A (CPA) among the top 100. This supports the high dependence of *Leishmania* amastigotes on papain-like cysteine peptidases in the mammalian host. In the spleen, tryparedoxin 1, tryparedoxin 2 and trypanothione reductase were amongst the top 100 most highly abundant mRNAs, suggesting that in the spleen, amastigotes adapt their transcriptional response to deal with an increase in environmental oxidative stress. This is in line with a previous study comparing a virulent *L. donovani* strain from Sri-Lanka with its avirulent counterpart that causes cutaneous leishmaniasis in which increases in stress response proteins and antioxidants were associated with changes in virulence and survival in visceral organs (37). On the other hand, we did not observe the expression of the protein kinases (LdBPK361580 and LdBPK361590) associated to the visceral-virulent phenotype in a recent genomic study in the same visceral vs cutaneous model (38).

Only 310 parasite genes out of the 4,998 expressed in the liver showed cumulative counts above 50 for both species, suggesting that low expressed genes would not be captured within this data, and statistically significant differences may not bear true biological differences when few reads are available as between group comparisons. This can result in exaggerated significance values and larger log2FC for lowly expressed genes with few counts assigned to the gene. This was not the case for the SPPEGs we identified. Of note, there was no expression of the double-strand break repair Rad21, reported as having increased expression in amastigotes by comparative transcriptomics of *L. infantum* and *L. major* (39). In addition, out of 8 protein kinases (PKAC1, MPK15, MPK10, EF2A2, PK4, CK2A1, CRK8, TOR3) recently identified as specific for amastigote infection and key for the survival of *L. mexicana* in the mouse (40), there was no expression detected for MPK15 and PK4 in the liver or spleen amastigotes, while the expression of PKAC1 and, EF2A2 was detected solely in spleen amastigotes. The expression of the parasite Inhibitor of Serine Peptidase (ISP2) was not detected (zero counts) in either liver or spleen amastigotes, in agreement with the prediction that its expression disfavours parasite development of *L. donovani* in macrophages (41).

In a recent dual-seq analysis of murine macrophages infected *in vitro* with virulent and avirulent *L. donovani* strains, the authors reported 96 parasite genes which were DE between the virulent and avirulent strains, with significant increases in proline oxidase, protein kinases and calpain-like peptidases in virulent parasites (14). The expression of the calpain-like gene (LDBPK_310430), reported by these authors as increased in virulent *L. donovani*, was below our cut-off for detection of expression in liver or spleen amastigotes. However, we observed high expression of another calpain-like gene (LdBPK_270510, within top 200) by parasites in the spleen and in the liver, which likely reflects heterogeneity among parasite strains regarding the expression of the individual members of the large family of calpain-like genes. In contrast, the authors identified tryparedoxin as upregulated in amastigotes of the avirulent strain as compared to their virulent strain, which is discrepant with our findings of the high tryparedoxin mRNA abundance by amastigotes in the tissues of infected mice.

We found only 4 SSPEGs between *L. donovani* and *L. infantum*, two of which were hypotheticals. The orthologue of the Lmsp7, a SSPEG expressed by *L. infantum*, is identified as a snRNA binding protein in *Trypanosoma sp*., and might represent a hub in the regulation of expression of other genes at the post-transcriptional level. Another SSPEG with increased expression in *L. infantum* is a ubiquitous enzyme involved in the generation of unsaturated fatty acids, including linoleic acid. Of relevance, exogenous linoleic acid was reported to decrease *L. donovani* survival in infected macrophages (42) and to enhance the inflammatory response of macrophages infected with *L. donovani* through LPO (43). However, we do not know if or how altered levels of unsaturated lipids in the parasite itself could affect infection.

In general, we found discrepancies between the expression of parasite genes previously associated with parasite virulence in transcriptomic studies in models of macrophage infections *in vitro* and the expression in amastigotes in the tissues of infected mice described here. This highlights the importance of care when extrapolating data from *in vitro* studies to an *in vivo* setting and by extension potentially limits the extrapolations of our findings in the mouse to human disease. A great deal of data on the immune response to visceral *Leishmania* species are available in mouse models of infection, especially in those employing the use of knock-out mice. Although the kinetics of the expression of cytokines and chemokines was previously addressed in BALB/c mice infected with *L. donovani* (17), a comparative paired analysis with *L. infantum* infections was not available. Here, we found most of the immune-related genes reported as upregulated at day 32 for mice infected with *L. donovani*, in both infections, denoting that the host responds to both parasite species in a similar manner. Adding to this picture, we observed significant upregulation of IL-27. The anti-inflammatory properties of IL-27 have been implicated in the prevention of protective Th1 responses to *L. donovani* and *L. infantum*, while limiting tissue damage (44). A linked cooperative action between IL-27 and IL-10 as a deactivating immunomodulatory arm during experimental visceral infection was recently proposed (45). In this study, IL-27 was responsible for reducing the levels of TNF and IFNγ required for parasite control, therefore favouring infection. However, we found abundant mRNA for *Tnf* and *Ifng* in the livers of infected mice, despite abundant *Il27* and *Il10* mRNA, suggesting that in the liver, IL-10 and IL-27 may be insufficient to attenuate the inflammatory response associated with parasite control. Considering that the relative amounts of cytokines/chemokines in tissues, as well as of their cell surface receptors, are highly susceptible to post-translational control by proteolysis, for example, our findings for the mRNA expression levels might not fully reflect the true nature of the tissue environment. Indeed, proteases known to inactivate cytokines or to cleave surface receptors, such as neutrophil elastase, were DEG in the infected organs.

Among the unique species-specific host DEGs, two might further impact the tissue microenvironment in ways that cannot not be verified by RNA-seq. Arachidonate 5-lipoxygenase, which was uniquely upregulated in the *L. donovani*-infected spleen, plays a dual role in inflammation due to its ability to enhance or to decrease the levels of lipid mediators, eicosanoids, and therefore the levels of pro-versus anti-inflammatory mediators that cannot be inferred based on gene expression alone. Furthermore, this enzyme can act as a hub for many cellular downstream responses related to migration and maturation, because it participates in dendritic cell migration, wound healing and monocyte adhesion to the endothelium. Changes in cellular migration and adhesion, for instance, cannot be addressed by RNA-seq. On the other hand, aldehyde oxygenase 1 (AOX1), uniquely upregulated in the *L. infantum*-infected spleen, and highlighted in the network analyses, has a broad oxidase activity, and can be a prominent source of superoxide generation, as well as contribute to nitric oxide (NO) production. Those findings reveal a greater chance for an enhanced oxidized environment in *L. infantum* infection, which could in turn contribute to the inactivation of certain cytokines, peptidase inhibitors and other players in tissue remodelling. We did not address potential nuances in tissue pathology, which requires detailed histological analyses.

Transcriptomics analyses in the hamster model of infection have identified mRNAs for regulatory cytokines (*IL4, IL10* and *IL21*), arginase (*Arg1*) and low nitric oxide production by macrophages as key factors of immunopathology and parasite persistence (19, 26). In the mouse, we observed significant increases in *IL10* and *IL21* mRNA accumulation in the spleens of infected mice, but differently from what seen in hamsters (26), *Nos2* was likewise highly increased in mice, whereas *Arg1* and *Arg2* (log2FC0.9, FDR=0.5) were not altered compared to uninfected control, despite high parasite loads. This highlights a major difference between the mouse and hamster models, and suggests that in the mouse, additional mechanisms beyond the nitric oxide-arginase axis may be pivotal to allow disease progression in the spleen. In common with the hamster studies, we observed enrichment in IFN-associated pathways, which were shown to unexpectedly promote parasite burden in *ex-vivo* splenic hamster macrophages. Also, in common with the hamster studies, we found a highly inflammatory pattern of gene expression, which contrasts with *in vitro* studies with murine or human macrophages that report mainly suppressive effects induced by the parasite. This highlights that an inflammatory environment in the tissue could have a profound effect in splenic macrophages. Although in hamsters infected with *L. donovani, IDO*-1 expression was highly upregulated both in the whole spleen tissue and in isolated splenic macrophages (19), we found only a modest increase in *IDO*-1 in the *L. donovani*-infected mouse spleen (log2FC2.2, FDR<0.01) while no significant induction was observed in the *L. infantum*-infected spleen (log2FC1.62, FDR<0.1). The enrichment in the tryptophan pathway found specifically for *L. donovani*-infected spleens is consistent with increased *IDO*-1 and with the kynurenine (KYNU) pathway for tryptophan catabolism. Adding to this picture, enzymes for phenylalanine and tyrosine metabolism were likewise among the *L. donovani*-induced unique DEGs, revealing an enrichment of genes related to nutritional metabolism. Intriguingly, such changes in *IDO*-1 and nutritional metabolism are also found in leprosy (36) and we have recently described at the cellular level, a close parallel of gene expression in macrophages infected *in vitro* with *L. donovani* (46) with those described for macrophages infected with *M. leprae* (47), including the induction of Type-I IFNs and OASL2.

The pathophysiology of bleeding has been attributed to disseminated intravascular coagulation (DIC). DIC is a disorder of the fibrinolytic and hemostatic systems that manifests as the generation of numerous and widespread microthrombi in the vasculature. Due to the increased spread and lack of compensatory control, DIC leads to eventual exhaustion of these pathways, resulting in propensity for diffuse bleeding and end organ complications. DIC is associated with numerous clinical diseases where excess cytokine production is observed, notably sepsis, and it has been suggested that VL in Brazil represents a slow to develop systemic inflammatory syndrome. Although the mouse model is not suited to address bleeding phenotypes nor DIC in the context of *Leishmania* infections, there was a significant enrichment of pathways related to coagulation and haemostasis in the infected animals, both in the liver and spleen. This is compatible with the major changes in the vascular architecture of those organs in infected animals and might reflect the host attempts to minimise tissue damage caused by excessive inflammation and coagulopathies. In the spleen, there was significant enrichment of pathways in both directions of clotting, i.e., formation and dissolution of fibrin clots, which may account for a better control of a bleeding phenotype in the mouse. A more robust response to visceralising *Leishmania* towards an equilibrium of hemostasis might be a key feature distinguishing the mouse model from human infections. Of note, several serpin genes (9 genes), which encode serine protease inhibitors, that inactivate enzymes of the coagulation cascade, including thrombin (48), and act on tissue remodelling (49), were significantly upregulated in infected mice. Serpins are also associated with attenuation of inflammation independently of their protease inhibitory properties (50). The role of microbial serpins in the control of hemorrhage and inflammation in a mouse model of lethal viral sepsis associated to DIC was recently reported, where clot inhibitors had moderate effects (51–53). In addition, *SERPIN3G*, which is implicated in the control of inflammation is highly upregulated (log2FC4). In conjunction, higher accumulation of serpins could contribute to ensure haemostasis and to attenuate inflammation in response to infection.

In summary, we provide a comprehensive paired dataset of parasite and host genes associated with infections with *L. donovani* or *L. infantum* in the mouse model. Although the host responded to both infections broadly in the same way, in the liver and spleen, we identified hundreds of host genes which were specifically DE in infections with either parasite species, which can serve as a source to address specific pathways of interest.

## Methods

### Ethics statement

Animal procedures were performed in accordance with and approved by the Ethics Committee on Animal Use of UFRJ (Comissão de Ética no Uso de Animais, CEUA) 034/15– UFRJ. Experiments were performed in accordance with the guidelines and regulations of the National Council of the Control of Animal Experimentation (Conselho Nacional de Controle de Experimentação Animal, CONCEA).

### Animals and parasites

Ethiopian *L. donovani* LV9 (MHOM/ET/67/HU3) and Brazilian *L. infantum* NLC (MHOM/BR/2005/NLC) were chosen for this study due to their geographical and phylogenetic divergence (54, 55). Indeed, Brazilian *L. infantum* isolates have been shown to have much lower nucleotide diversity compared to Indian and African *L. donovani* (55). The parasites were grown as promastigotes in modified Eagle’s medium, designated HOMEM (Thermo Fisher Scientific, Waltham, MA, USA), that was supplemented with 10% heat-inactivated fetal bovine serum (FBS, Thermo Fisher Scientific) and incubated at 25 °C. Amastigotes of *L. donovani* and *L. infantum* were purified from the spleens of Syrian hamsters that were infected 4 months previously by i.p inoculation of 3 × 10^7^ amastigotes. To harvest parasites for murine infections, spleens from infected hamsters were macerated through a 70 µm cell strainer (BD Biosciences, San Jose, CA, USA) in Balanced Salt Solution (BSS), centrifuged at 300 *g* for 5 min, the supernatant was collected and centrifuged again at 2,600 *g* for 10 min. The cell pellet was then washed in BSS at 2,600 *g* for 10 min and counted. 3 × 10^7^ amastigotes in 100 µl PBS were injected into a single cohort of 15 BALB/c mice. Animals were monitored daily for any unexpected adverse clinical symptoms (n=5 per group).

### RNA extraction, purification and cDNA reverse transcription

Two pieces of each organ (5 mg) were cut and kept in RNAlater (Thermo Fisher) until further processing. Tissues were thawed on ice and immediately transferred to 0.6 mL lysis buffer (PureLink RNA Mini kit, Thermo Fisher, USA). Tissues were homogenized using plastic pestle homogenizer for microtubes for about 5 min, after homogenization RNA extraction was performed using the PureLink RNA Mini kit, according to the manufacturer’s instructions. Total RNA was eluted in RNAse-free water and quantified using Qubit™ RNA HS Assay Kit (Thermo Fisher, USA, #Q32852). RNA integrity was assessed by non-denaturing 1.2 % (w/v) agarose electrophoresis. Genomic DNA was removed by treatment with Ambion DNA-free kit according to standard manufacturer’s instructions. Five hundred nanograms to one microgram of total RNA were reverse transcribed into cDNA with SuperScript™ VILO™ Master Mix (Thermo Fisher, USA, #11755050) according to the manufacturer’s directions. The resulting samples were then prepared for sequencing using the NEB next II ultra kit. One mouse (M4) from the *L. infantum*-infected group and one mouse (M4) from the uninfected control group were removed from subsequent analysis due to poor read counts likely the result of technical error.

### Sequencing and bioinformatics analyses

Adaptors were first removed from reads using Cutadapt v. 1.8.3 with the parameters –a, -A and –p to remove universal adaptors from both 5’ and 3’ ends (56). Singletons were removed, and reads were further trimmed and scored for quality using sickle v. 1.33, using the options pe for paired end and options –f –r –t sanger for base quality scoring method, and –q 20 and – l 20 to set discard read length and quality score filter based on a phred score (57). Reads were then aligned using STAR aligner v. 2.5.1b using–sjdbOverhang 150. Indexes were generated through the concatenation of annotations from TritrypDB of *L. infantum* and *L. donovani* v46, with the GRCm38.p6 ENSEMBL annotation of *Mus Musculus*, and reference fastas (58–60). Alignments were run using these combined parasite and mouse transcriptome references using –--outFilterMatchNmin 40 to locally align to account for splice leader sequences within parasite transcripts, which may otherwise be discarded. Reads from individuals from both groups were aligned to the same reference to reduce the identification of false positive SSPGs due to differences in gene length between *L. donovani* and *L. infantum* orthologues. Reciprocal alignments were done, and DE lists from both were used. The IDs of SSPGs were compared using the syntenic orthologues from tritrypdb only, and paralog counts in parasite DE genes were investigated to see whether copy differences resulted in them being identified as DE, these are included in the **Table S2**.

HTseq-count was used with the union algorithm to assign reads to genes (61). EdgeR v 3.30.3 was used to identify DE genes and normalise and perform pairwise comparisons using a generalised linear model (62). Identification of parasite SSPE genes was done using alignments from the parasite only, due to the greater abundance of host transcripts, which otherwise mask the variability observed in the parasite transcripts. Baseline SSPE counts are unfiltered, filtered DE counts use a cut-off of log2FC1, which allows for validation by qPCR. edgeR was run using both counts per million (CPM) and transcripts per million (TPM) methods, where TPM controls for gene length to check whether SSPE genes were identified due to differences in gene length between *L. donovani* and *L. infantum* orthologues. RPKM values were also calculated using *L. infantum* infected individuals against the *L. infantum* reference, and the *L. donovani* infected individuals against the *L. donovani* reference, and log2FCs calculated between the groups.

PCA analysis was performed in R 4.0.3 using the princomp function to generate eigenvalues from the transcriptomes. Vst normalisation was used on the count matrix prior to eigenvalue conversion to aid clustering. Volcano plots were generated in native R (63). MA plots were derived using the maPlot function within the edgeR package using logCPM values for the log abundance and log2FC values calculated in edgeR (62, 63). The plots were not smoothed (smooth.scatter = FALSE) and the data was not fitted to a lowess curve. A smearwidth of 0.5 and a glmLRT method was used for DE identification.

### Pathway analysis

Gene symbol annotations were derived using the ENSEMBL biomart database (59). Host DE genes were assigned to their GO terms through STRINGDB’s annotation db v. 11.0 and enrichR [last accessed 13/4/21] (64, 65). Annotations for the *Leishmania* DE genes were obtained from tritrypdb (60).

Networks were identified using confidence to connect network edges, with line thickness indicating the strength of the association. This support was from text mining, experimental data, co-expression, database collation, gene fusion studies, neighbourhood (genomic location) and co-occurrence from expression studies. Networks were filtered through using confidence interaction values between 0.4-0.9. Disconnected nodes from DE lists were removed.

Msigdb was used within GSEA to identify overlaps between the hallmark genesets and the C7 immunology panel against genes that were commonly DE to both parasites in the spleen and liver, as well as genes uniquely DE for each parasite (66).

### Statistics and experimental design

As this was a comparative study not designed to identify any formal statistical significance between infection groups, no formal sample size calculation was performed. We used the same sample sizes per group (n=5) as those from our previous study, which were sufficient to identify gene signatures (17). Mice were infected as a single cohort for each experiment (as detailed in Results). Data from Fig. 1B, C was analysed and the figure constructed using GraphPad Prism 8 software (GraphPad Software, San Diego, CA). The remaining figures were generated in R v. 4.0.3 (63). After assessment for normality, data on tissue weights and parasite loads were analysed using the Mann Whitney test and a p-value of less than 0.05 was taken as significant. Statistical analysis and graphical presentation of transcriptomic data is described above.

## Data availability

Raw sequencing data and the gene count tables have been deposited and are publicly available from GEO Gene expression Omnibus database under the accession GSE143799.

## Acknowledgements

This work was funded by a National Centre for Replacement, Refinement and Reduction of Animal in Research (NC3Rs) / Innovate UK CRACKIT Challenge Award (NC.CO13117 and NC/CO13295 to PMK, and JM) and the Medical Research Council Newton (MR/M026167/1 and MR/N017269/1 to JCM, PK and APCAL). PMK was supported by a Wellcome Trust Senior Investigator Award (WT104726). APCAL was supported by FAPERJ (grant number E26/202.655/2019) and CNPq (grant number 311208/2017-7) and is a CNPq fellow. The funders had no role in study design, data collection and analysis, decision to publish, or preparation of the manuscript.

## Figure legends

**Fig. S1.**
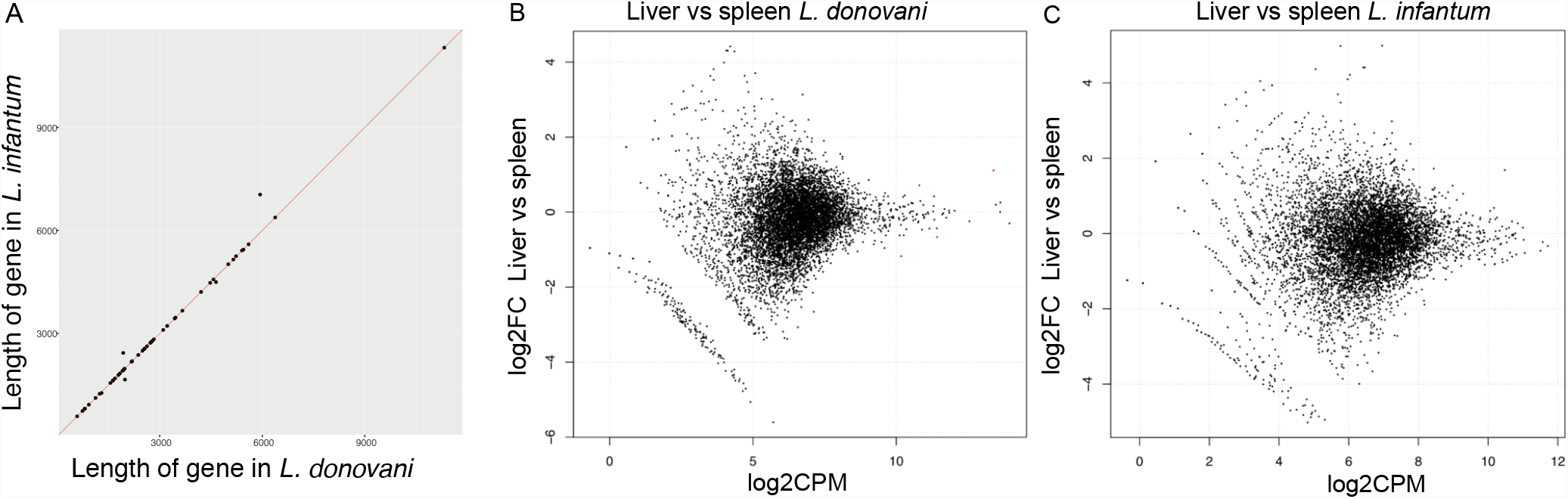
**(A)** is a plot of the gene length of genes that have a positive log2FC RPKM value and a negative log2FC CPM value. This shows the length of the syntenic orthologues used in *L. infantum* and *L. donovani*. **(B)** is an MA plot with the DE genes identified in parasite genes between *L. donovani* liver and spleen samples. **(C)** is an MA plot with the DE genes identified in parasite genes between *L. infantum* liver and spleen samples.

**Fig. S2.**
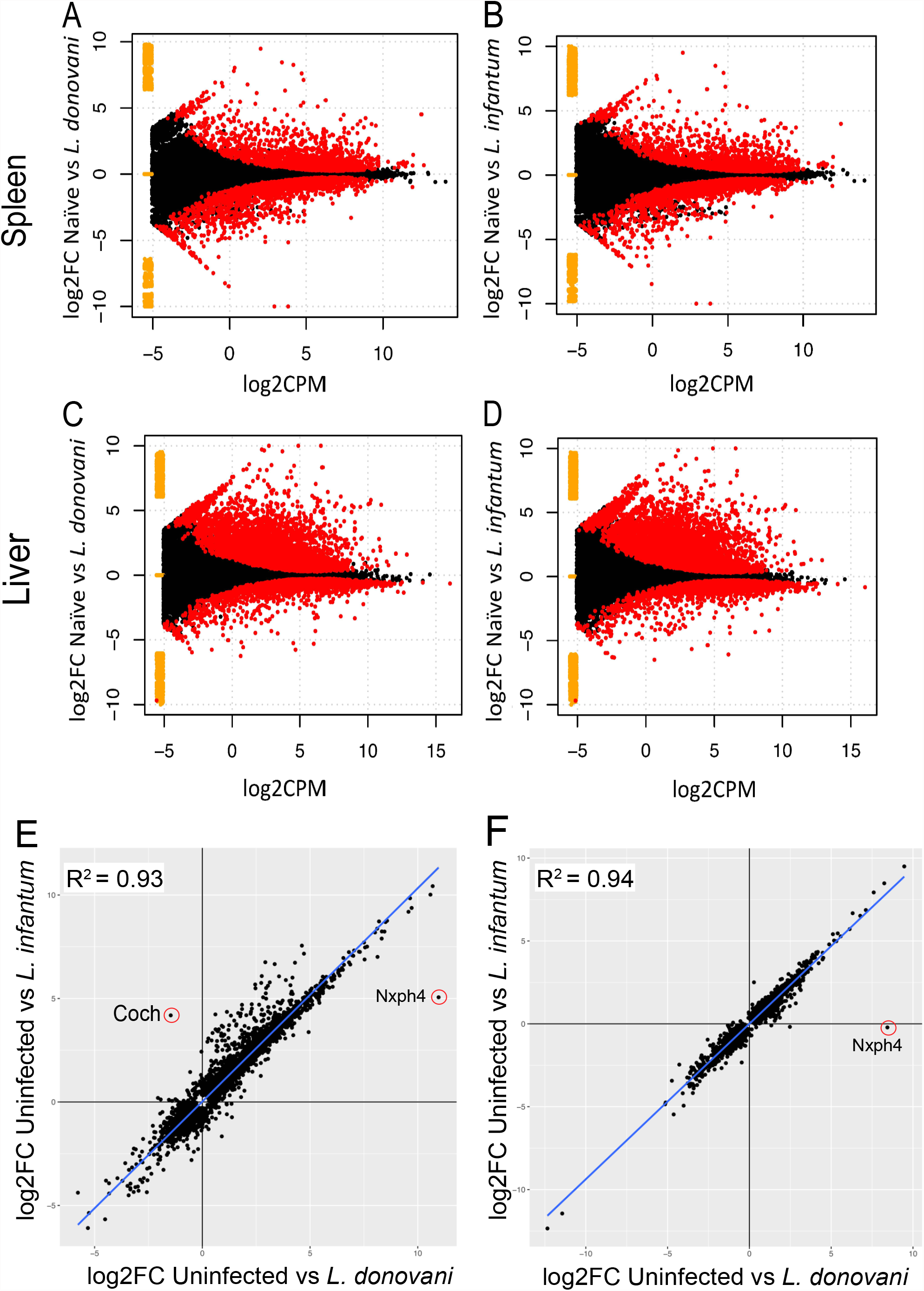
**(A)** is an MA plot with the DE genes identified in host genes between uninfected and *L. donovani* infected mice in the spleen **(B)**, uninfected and *L. infantum* infected mice in the spleen, DE shown in red. **(C and D)** show MA plots with the same comparisons, but in the liver. Yellow points represent genes expressed in one condition, but not the other. **(E)** Correlation between log2FC for each gene in the host genes in the liver for genes with a logCPM <1 removed, log2FC taken from **Table S3**. Line of best fit, and R^2^ displayed. **(F)** Correlation between log2FC for each gene in the host genes in the spleen.

**Fig. S3.**
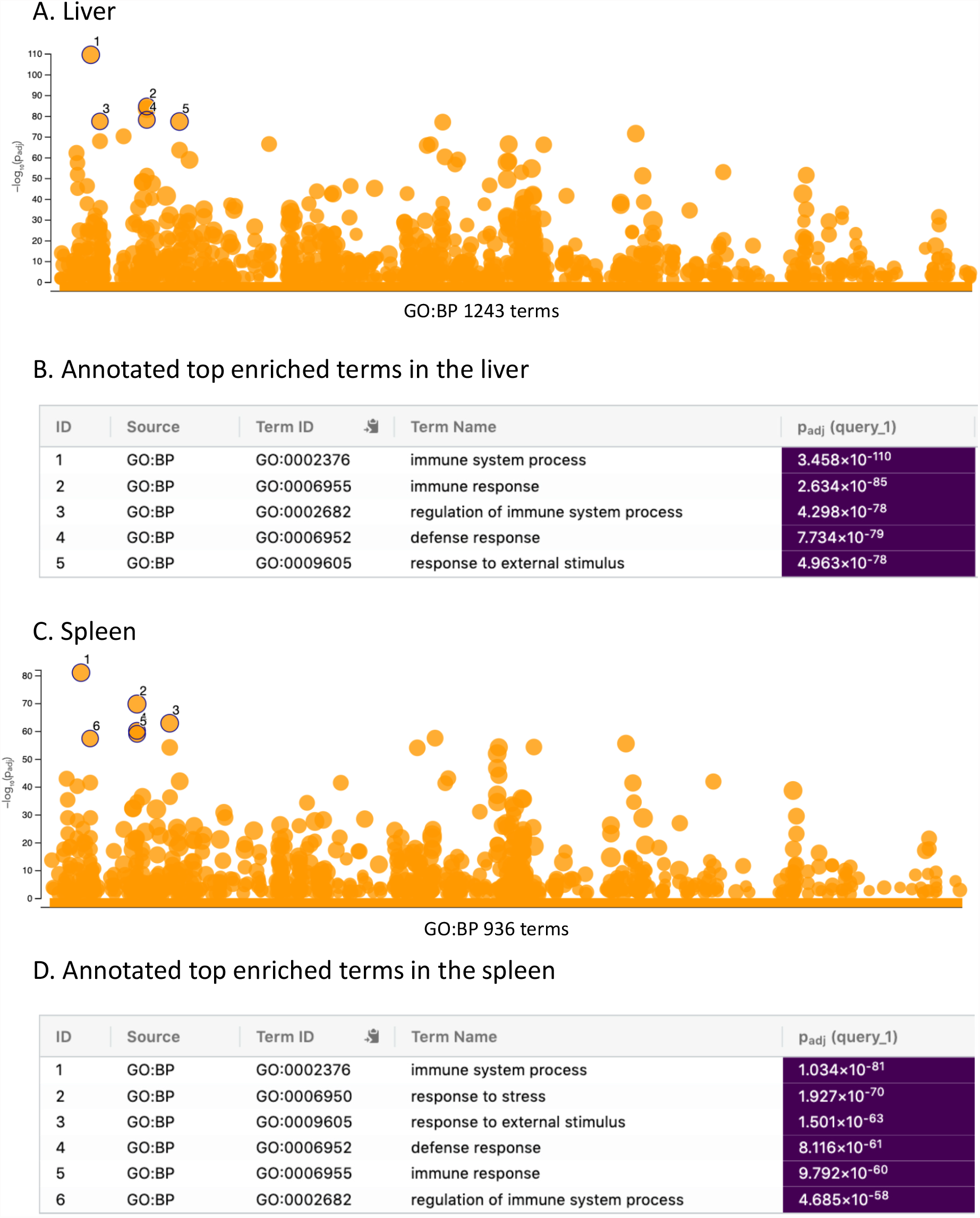
**(A)** G profiler results for GO Biological process enrichments. Each circle represents a term, and the adjusted p-value of the enrichment. The genes used in this enrichment are from all DE to both species relative to naïve in the liver. The top 5 most enriched terms are given in **(B)** and the significance of this enrichment. **(C)** shows this but for common DE in the spleen. **(D)** shows the top 6 enriched terms in the spleen because this contains the top 5 enriched terms in the liver, and an additional term which is the second most enriched term. Terms 4 and 5 overlap due to very similar log10 enrichment p values.

**Fig. S4.**
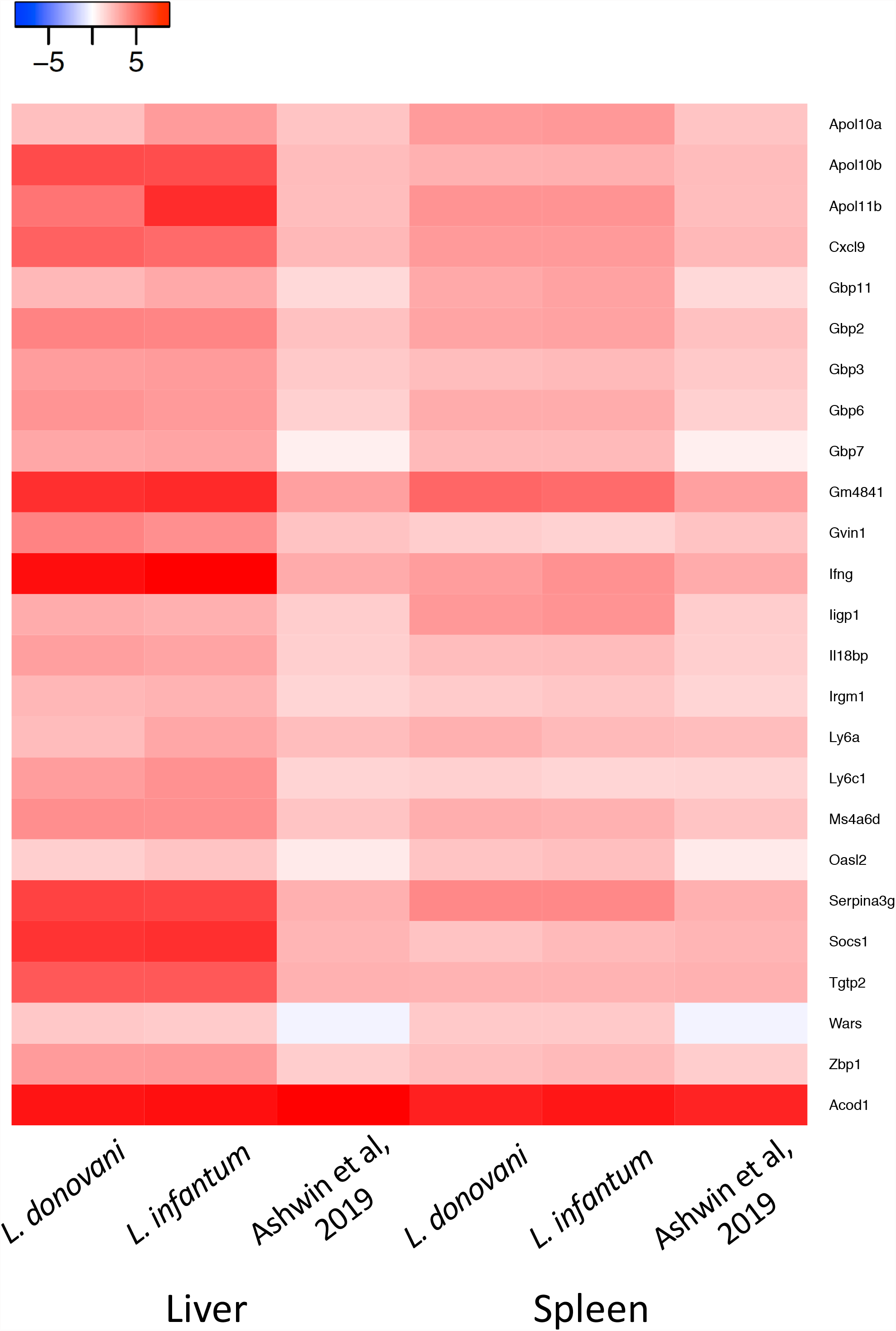
expression based on normalised counts for the 25 genes previously published as a 26 gene signature. These are shown for the liver for log2FC values in *L. donovani* vs naïve, then in *L. infantum*, and then the value compared to the microarray expression of these genes in (18) in the first three columns respectively, and in the same order for the spleen.

**Table S1** Raw counts and the log2FC, logCPM values for DE genes in the spleen, in the liver for the parasite genes. Spleen SSPEGs are the species-specific DE genes. Genes that were unexpressed in *L. donovani* are also given.

**Table S2** DE genes between *L. donovani* and *L. infantum* parasite genes in the spleen that had paralogous genes that had differing counts between *L. infantum* and *L. donovani* and their annotation.

**Table S3** Raw counts, log2FC, logCPM values and list of DE genes in the spleen and liver for the host genes. DE genes are given for all three pairwise combinations of uninfected *L. infantum* and *L. donovani* infected groups. Gene lists for unexpressed genes are available alongside **Table S3** on GEO under accession GSE143799, supplementary material GSE143799_Mouse_counts.xlsx

**Table S4** EnrichR enrichments for the host genes in the liver and list of host genes specifically DE in infections with either *L. donovani* (221) or *L. infantum* (429).

**Table S5** EnrichR enrichments for the host genes in the spleen and list of host genes specifically DE in infections with either *L. donovani* (286) or *L. infantum* (186).

**Table S6** Details of genes involved in the networks identified using STRINGDB used to generate networks in **Fig. 7**. String mapping sheets give the genes used and the mapping information for each per panel in **Fig. 7**. The enrichments shown in **Fig. 7** include the gene IDs of disconnected nodes not connected to the networks, but still used in the enrichments. The resulting networks are also shown for *L. donovani* specific, *L. infantum* specific and common gene networks in the spleen.

